# Shared and dissociable features of apathy and reward system dysfunction in bipolar I disorder and schizophrenia

**DOI:** 10.1101/546036

**Authors:** Matthias Kirschner, Flurin Cathomas, Andrei Manoliu, Benedikt Habermeyer, Joe J. Simon, Erich Seifritz, Philippe N. Tobler, Stefan Kaiser

**Affiliations:** Department of Psychiatry, Psychotherapy and Psychosomatics, Psychiatric Hospital, University of Zurich, 8032 Zurich, Switzerland; Montreal Neurological Institute, McGill University, Montreal, Quebec, Canada; Psychiatric Services Aargau, 5201 Brugg, Switzerland; Department of General Internal Medicine and Psychosomatics, Centre for Psychosocial Medicine, Heidelberg, Germany; Department of Psychosomatic Medicine and Psychotherapy, Medical Faculty, Heinrich-Heine-University Düsseldorf, Dusseldorf, Germany; Neuroscience Center Zurich, University of Zurich, 8057, Zurich, Switzerland; Zurich Center for Integrative Human Physiology, University of Zurich, 8057 Zurich, Switzerland; Laboratory for Social and Neural Systems Research, Department of Economics, University of Zurich, 8006, Zurich, Switzerland; Division of Adult Psychiatry, Department of Psychiatry, Geneva University Hospitals, Chemin du Petit-Bel-Air, 1225 Chêne-Bourg, Switzerland

**Keywords:** bipolar disorder, euthymic, apathy, diminished expression, fMRI, monetary incentive delay task, negative symptoms, reward anticipation, striatum, transdiagnostic

## Abstract

**Background:** Bipolar disorder I (BD-I) is defined by episodes of mania, depression, and euthymic states. These episodes are among other symptoms characterized by altered reward processing and negative symptoms (NS), in particular apathy. However, the neural correlates of these deficits are not well understood.

**Methods:** We first assessed the severity of negative symptoms in 25 euthymic BD-I patients compared to 25 healthy controls (HC) and 27 patients with schizophrenia (SZ). Then, we investigated ventral and dorsal striatal activation during reward anticipation in a Monetary Incentive Delayed Task and its association with NS.

**Results:** In BD-I patients NS were clearly present and the severity of apathy was comparable to SZ patients. Apathy scores in the BD-I group but not in the SZ group correlated with sub-syndromal depression scores. At the neural level, we found significant ventral and dorsal striatal activation in BD-I patients and no group differences with HC or SZ patients. In contrast to patients with SZ, apathy did not correlate with striatal activation during reward anticipation. Explorative whole brain analyses revealed reduced extra-striatal activation in BD-I patients compared to HC and an association between reduced activation of the inferior frontal gyrus and apathy.

**Conclusion:** This study found that in BD-I patients apathy is present to an extent comparable to schizophrenia, but is more strongly related to sub-syndromal depressive symptoms. The findings support the view of different pathophysiological mechanisms underlying apathy in the two disorders and suggest that extra-striatal dysfunction may contribute to impaired reward processing and apathy in BD-I.

## Introduction

Bipolar disorder I (BD-I) is a severe neuropsychiatric illness that causes extensive individual morbidity and socioeconomic burden (Vos *et al.*, 2012). Clinically, the disorder is defined by the occurrence of at least one manic episode and often includes recurrent states of depressive, manic, mixed and euthymic episodes (Association, 2013). It is increasingly becoming evident that processing of motivation and reward are affected in all of these states (Ashok *et al.*, 2017). This is supported by recent findings showing a transdiagnostic dopamine dysfunction spanning from bipolar disorder to schizophrenia (Jauhar *et al.*, 2017). In order to understand the relationship between dysfunctional motivational processes and clinical disease manifestations within but also between different diagnostic entities, valid dimensional psychopathological constructs are essential (Cuthbert and Insel, 2013).

Negative symptoms (NS) have been proposed as such a construct (Strauss *et al.*, 2016). Originally considered as a hallmark symptom complex of schizophrenia (Kraepelin, 1921), it has become evident that NS also occur outside of the schizophrenia-spectrum (Strauss and Cohen, 2017). A recent study reported that NS can indeed be measured in BD-I patients (Strauss *et al.,* 2016). NS can be grouped into a motivational dimension (hereafter referred to as apathy) containing anhedonia, avolition and asociality, and a diminished expression dimension, which includes blunted affect and alogia (Blanchard and Cohen, 2006, Strauss *et al.*, 2013). Critically, increasing evidence from several fields suggests that the two dimensions are caused by different neurobiological and behavioural mechanisms (Caravaggio *et al.*, 2018, Cathomas *et al.*, 2015, Hager *et al.*, 2015, Hartmann *et al.*, 2015, Wolf *et al.*, 2014). This differentiation might therefore be critical for the development of effective treatment (Galderisi *et al.*, 2018, Kaiser *et al.*, 2017). With respect to the relevance of NS in BD several studies have reported elevated NS scores even in euthymic patients (Hawkins *et al.*, 1997, Mancuso *et al.*, 2015, Strauss *et al.*, 2016), which are strongly associated with functional impairments (Atre-Vaidya *et al.*, 1998, Samalin *et al.*, 2016, Serafini *et al.*, 2018). Studies comparing the severity of negative symptoms between SZ and BD yield mixed results (Strauss and Cohen, 2017, Tso *et al.*, 2014) but suggest that NS might be more stable and trait like in SZ. With respect to the two negative symptom dimensions a recent study found that BD patients only differed from SZ patients on the diminished expression dimension but showed similar severity of apathy (Strauss *et al.*, 2016). Furthermore, several studies have reported residual anhedonic symptoms in euthymic BD patients, which were comparable to SZ patients (Di Nicola *et al.*, 2013, Tso *et al.*, 2014) and relevant for impaired reward learning (Pizzagalli *et al.*, 2008). Although comparable in trait anhedonia, SZ patients showed more severe total negative symptoms, lower experience of pleasure and lower behavioral activation (Tso *et al.*, 2014).

It is well established that striatal dysfunction during reward anticipation plays an important role in the pathophysiology of NS in patients with schizophrenia (SZ) (Heinz and Schlagenhauf, 2010, Nielsen *et al.*, 2018, Radua *et al.*, 2015). Recent studies suggest a specific link between reduced ventral (VS) and dorsal striatal (DS) activity and apathy (Kirschner *et al.*, 2016b, Mucci *et al.*, 2015, Simon *et al.*, 2010, Stepien *et al.*, 2018, Wolf *et al.*, 2014). In contrast, results regarding reward anticipation in euthymic BD patients are conflicting, with studies showing both increased (Mason *et al.*, 2014, Nusslock *et al.*, 2012) and unaltered (Caseras *et al.*, 2013, Dutra *et al.*, 2015) VS activity compared to healthy controls (HC). To our knowledge the association between NS and striatal dysfunction has not been investigated in patients with BD-I. A transdiagnostic approach taking advantage of the current findings from schizophrenia research provides a unique opportunity to elucidate the neural substrates of motivational deficits in BD. Furthermore, given that only recently extra-striatal prefrontal correlates of apathy have been observed (Dowd *et al.*, 2016, Wang *et al.*, 2016), this transdiagnostic approach allows the comparison of potential contributions of extra-striatal dysfunction to the formation of apathy in BD and SZ.

The aims of the present study therefore were 1) to explore, on a psychopathological level, the association between apathy and other symptom dimensions in BD-I patients currently in an euthymic state compared to patients with SZ (Kirschner *et al.*, 2016b), 2) to investigate whether euthymic BD-I patients show impaired VS or DS activity during reward anticipation during a variant of the Monetary Incentive Delay (MID) task, 3) to compare BD-I patients to patients with SZ regarding the association between VS or DS activity during reward anticipation and the two negative symptom domains apathy and diminished expression and 4) to study extra-striatal differences between BD-I patients and HC using an explorative whole brain analysis approach.

## Material and Methods

### Participants

25 participants with a diagnosis of BD-I and 27 patients with a diagnosis of schizophrenia (Kirschner *et al.*, 2016b) were recruited from in- and outpatient units of the Psychiatric Hospital of the University of Zurich, Switzerland and affiliated institutions. 25 HC, all currently employed, were recruited from the general community. For all participants the inclusion criterion for age was 18 to 55 years. Regarding the recruitment of euthymic BD-I patients, we included 18 patients from outpatient units and 7 patients from inpatient units, which were at the end of their hospitalization. The data from SZ patients (11 outpatients and 16 inpatients) and HC rely on our previously published study (Kirschner *et al.*, 2016b).

All patients had a stable medication for at least 14 days and a dose of lorazepam not exceeding 1mg daily. Using the structured Mini-International Neuropsychiatric Interview for DSM-IV (Ackenheil *et al.*, 1999), we A) confirmed the diagnosis of BD-I or schizophrenia, B) excluded patients with a current manic, hypomanic or major depressive episode, C) excluded patients with a history of schizoaffective disorder, and D) excluded patients with any other DSM-IV Axis I disorder. Healthy controls were screened for neuropsychiatric disorders using the structured Mini-International Neuropsychiatric Interview to ensure that they had no previous or present psychiatric illnesses. All participants were required to have a normal physical and neurologic status (including the assessment of extrapyramidal side effects, i.e. a total score ≤2 on the Modified Simpson-Angus Scale [MSAS] (Simpson and Angus, 1970) and no history of major head injury or neurological disorder. Patients with BD-I were clinically euthymic and on a stable medication for at least two weeks before the study. None of the patients with BD-I showed more than sub-syndromal depressive symptoms (Hamilton Depression Rating Scale, HAMD 17 Score < 17), as defined by the International Society for Bipolar Disorder (ISBD) Task Force (Tohen et al. 2009). The absence of manic symptoms was confirmed with the Young Mania Rating Scale (Young *et al.*, 1978) (mean= 0.4, SD = 0.91, min = 0, max = 4). The study was approved by the local ethics committee of the canton of Zurich and all participants provided written informed consent.

### Clinical and neuropsychological assessment

NS were assessed using the Brief Negative Symptom Scale (Kirkpatrick *et al.*, 2011) to answer all research questions in the present study. For comparison with earlier studies, additional assessment of NS using the Scale for the Assessment of Negative Symptoms (SANS) (Andreasen NC. Scale for the Assessment of Negative Symptoms (SANS). Iowa City, IA: University of Iowa; 1983) can be found in the Supplementary Results. Apathy was defined based on the BNSS subscales anhedonia (items 1-3), asociality (items 5, 6) and avolition (items 7, 8) and diminished expression was defined as the sum of the BNSS subscales blunted affect (items 9-11) and alogia (items 12, 13) (Kirkpatrick *et al.*, 2011, Mucci *et al.*, 2015). Please note, that the BNSS total score also includes the BNSS distress item, which was not included in one of the two factors (Kirkpatrick *et al.*, 2011, Mucci *et al.*, 2015).

Additional psychopathological assessment included the Hamilton Depression Rating Scale 21 (Hamilton, 1967) and Calgary Depression Scale for Schizophrenia (CDSS) (Addington *et al.*, 1990) to assess depressive symptoms. The CDSS has also been validated for patients with major depressive disorders (Micoulaud-Franchi *et al.*, 2018) and has been suggested as transdiagnostic tool to assess subclinical depressive symptoms across BD-I patients and SZ patients. The Global Assessment of Functioning Scale (GAF) (Frances et al., 1994) and Personal and Social Performance Scale (PSP) (Juckel et al., 2008) were used to assess global level of functioning and the Positive and Negative Syndrome Scale (PANSS) (Kay et al., 1989) was used to assess all psychotic symptom dimensions. The PANSS factor scores were calculated according to the five-factor model of Wallwork et al. (Wallwork *et al.*, 2012). Since the different factor scores do not include all the PANSS items, sums of the subscales do not add up to the total PANSS score. All participants performed a comprehensive neuropsychological test battery, which has been used in previous studies (Hager *et al.*, 2015, Hartmann *et al.*, 2015) (Supplementary Methods).

### MID task

We employed a variant of the MID (Knutson *et al.*, 2000) with stimuli based on the Cued-Reinforcement Reaction Time Task (Cools *et al.*, 2005). This modified version was originally developed by Simon and colleagues (Simon *et al.*, 2015) and used in previous studies to investigate reward anticipation (Kirschner *et al.*, 2016b, Kirschner *et al.*, 2018, Simon *et al.*, 2015). In this variant, reward amount was directly determined by the individual response time of each participant (Fig S2). This adaptation allowed us to investigate the motivational properties of reward anticipation in the presence of strong action-outcome contingencies. Briefly, before starting the experiment participants were informed that they would receive all the money won during the two experimental sessions. At the beginning of each trial, one of three different cues was presented for 0.75 s. The cue indicated the maximum amount participants could gain (i.e., 2 Swiss francs (CHF), 0.40 CHF, or 0 CHF; 1 CHF = $1.03 US). After a delay of 2.5–3 s, participants had to identify an outlier from 3 presented circles and press a button as fast as possible. Immediately after the button press, participants were notified of the money they had won (duration of feedback 2 s) (Fig. S1). Error trials were defined as trials with an incorrect or late response (after 1 s). In all other trials, we calculated the pay-out structure for each trial on the basis of the response times of the previous 15 individual trials (Fig. S2). Therefore, the amount of money won depended on the response time in the current trial in relation to the response time in the previous 15 trials. The maximum amount of money to be won was 50 CHF. Every participant performed two training runs, one outside and one inside the scanner. Excluding the training sessions, the experiment included two runs with 36 trials of about 10 s each. The inter-trial interval (ITI) was jittered from 1 to 9 s with a mean of 3.5 s to enhance statistical power. In total, 1 run lasted about 6 min.

### Functional image acquisition

We used the same protocol as in our previous studies (Kirschner *et al.*, 2016a, Kirschner *et al.*, 2016b), which is described in the Supplementary Methods.

### Data analyses

Data (demographic, clinical, neuropsychological and behavioural) were analysed using SPSS (version 23, SPSS Inc.) and test for normal distribution was applied for all data. Group differences in sociodemographic and neuropsychological characteristics were investigated using 2-sample t tests for continuous and χ2 tests for categorical data. For non-normally distributed data, Mann–Whitney U tests were applied. The association symptom dimensions were investigated using bivariate Spearman correlations (r_s_). In particular, apathy was correlated with potential secondary sources for NS including depressive symptoms, positive symptoms and antipsychotic dose equivalents. FMRI data were analysed using SPM8 (revision 5236, <u>04-Feb-2013</u>) (Statistical Parametric Mapping, Wellcome Trust Centre for Neuroimaging, London, UK). The statistical tests were selected for separate comparisons of BD-I patients with HC or SZ, excluding the previously reported comparison of HC with SZ (Kirschner *et al.*, 2016b).

### Behavioural data analyses

The main behavioural outcome measure was response time (RT), defined as the time between target presentation and pressing the correct button. We conducted a 2-way repeated measures analysis of variance (ANOVA) with RT as dependent variable, group as between-subjects factor and reward condition (neutral, low, high) as within-subject factor. We performed the Mauchly test for the assumption of sphericity and in case of violations report Greenhouse–Geisser–corrected degrees of freedom. Bonferroni-corrected pairwise comparisons were calculated as *post hoc* tests for significant main effects. We performed correlation analysis between negative symptom factors and reward-related speeding. Reward-related speeding was calculated by subtracting the RT during the neutral condition (CHF 0) from the RT during the high reward condition (CHF 2.0). One BD-I subject was excluded only from the behavioural analysis due to corrupted data files. To account for multiple comparisons (group comparison of BD-I patients with HC and patients with SZ separately), we considered findings at p<0.025 as significant.

### Image pre-processing

The Image pre-processing followed the same protocol as in our previous studies (Kirschner *et al.*, 2016a, Kirschner *et al.*, 2016b) and is outlined in the Supplementary Methods.

### First- and second-level image analyses

A general linear model (GLM) approach was applied to assess data in an event-related design at the first level. For the three different reward anticipation phases, separate regressors were included: anticipation of no reward (CHF 0), anticipation of low reward (CHF 0.40) and anticipation of high reward (CHF 2.0). For the outcome phases, regressor for each condition (three basic regressors) was included. Additionally, for the low and high reward conditions the two outcome regressors were parametrically modulated by the actual outcome amount of each trial. Target presentation (one regressor) and anticipation, target and outcome phase in error trials (three regressors) were modelled as regressors of no interest. In total, the first-level model included 12 regressors. The canonical hemodynamic response function was used for convolving all explanatory variables. For reward anticipation the contrast anticipation of high reward (CHF 2.0) versus anticipation of no reward (CHF 0) was calculated. At the second-level analysis, individual contrast images of all participants were included in a random-effects model. Within-group activation was calculated using a 1-sample *t* test and between-group activation using a 2-sample t test. For all whole brain analysis the statistical threshold was set to p<0.05, whole-brain cluster-level family-wise error (FWE) rate corrected for multiple comparisons with a cluster-defining voxel-level threshold of p<0.001 uncorrected.

### Region of interest image analysis

The VS and DS were defined as regions of interest (ROIs) during anticipation of reward. We derived VS coordinates (Montreal Neurological Institute [MNI]) from a meta-analysis (Knutson and Greer, 2008) (left: x= −12, y= 10, z= −2; right: x= 10, y= 8, z= 0; 9mm sphere in one single VS ROI). DS coordinates were derived from a previously published fMRI data, indicating DS activation in response to the MID task (left: x=-9, y=3, z=15; right: x=9, y=3, z=15; 9mm sphere in one single DS ROI) (Yip *et al.*, 2015). This ROI approach was adapted from (Yip *et al.*, 2015) and has been used in our previous studies (Kirschner *et al.*, 2016a, Kirschner *et al.*, 2016b). The ROIs were constructed with the Wake Forest University Toolbox implemented in SPM8 (Maldjian *et al.*, 2003). The statistical thresholds at the voxel level were set to a family-wise error (FWE) corrected threshold for multiple comparison of p = 0.05 in each ROI. Mean percent signal changes were extracted for all voxels in the VS and DS using the REX toolbox (http://web.mit.edu/swg/software.htm). In other words, extraction and average calculation was not restricted to the significant voxels but to all voxels of the ROIs. In addition, we performed an extra-striatal ROI analysis in the prefrontal cortex, which was based on previous findings showing an association between a behavioural measure of apathy and reduced inferior frontal gyrus (IFG) activation (Kluge *et al.*, 2018). Structural masks of the left and right IFG were derived from the IBASPM 71 atlas implemented in the Wake Forest University Toolbox in SPM8 (Maldjian et al., 2003).

### Correlation analysis

The main hypothesis was tested by calculating bivariate Spearman correlations (r_s_) between negative symptoms (apathy and diminished expression) and percent signal change in the VS and DS. Finally, the Steiger test for dependent correlation coefficients was performed to test for potential differences between these correlations (Steiger, 1980).

## Results

### Sample characteristics

Participant characteristics, clinical data and group comparisons are summarized in Table 1. Patients with BD-I had a significant higher average number of education years compared to HC and SZ patients. Patients in the BD-I group were older than SZ patients and were characterized by a longer duration of illness compared to the SZ group. On both the Global Assessment of Functioning (*t* = – 2.613, *p* = 0.012) and the Personal and Social Performance Scale *(t* = −2.163, *p* = 0.036) BD-I patients scored higher, indicating a higher level of functioning compared to patients with SZ. In the BD group 18 out of 25 patients were treated with atypical antipsychotics, 18 patients received mood stabilizer and 7 patients antidepressants. All patients with SZ received atypical antipsychotics. Mean chlorpromazine equivalents were higher in patients with SZ compared to BD-I patients (BD-I: 185.99 ± 259.8, SZ: 508.01 ± 369.2; U = 133, p < 0.001). None of the patients received typical antipsychotics.

**Table 1.**
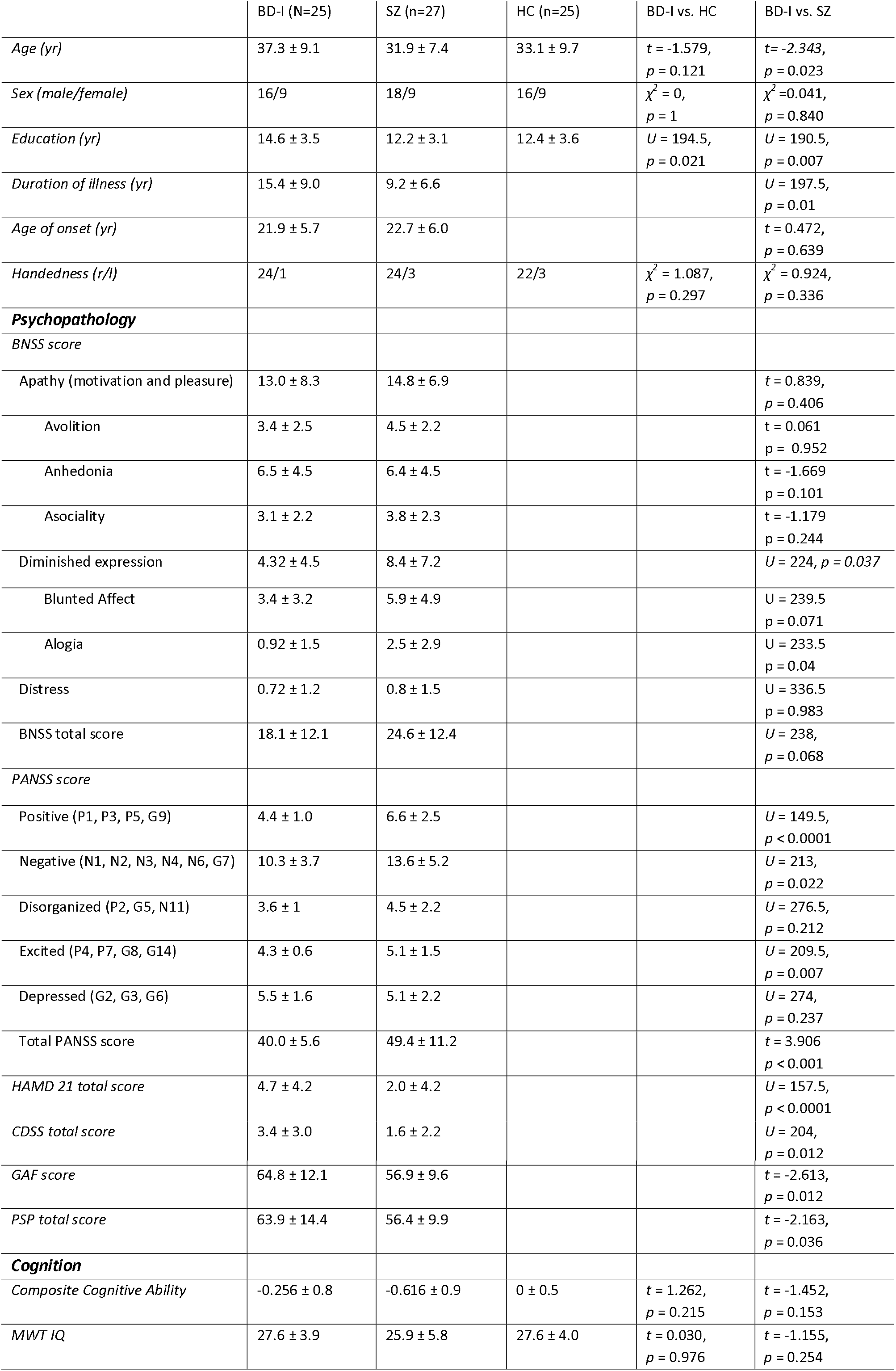
Sample Characteristics. Summary of the socio-demographic and neuropsychological characteristics of the study participants. Data are presented as average ± standard deviation. Abbreviations: BD-I: Bipolar Disorder I; SZ: Schizophrenia; HC: Healthy controls; BNSS: Brief Negative Symptom Scale. BNSS Apathy was defined based on the BNSS subscales anhedonia (items 1-3), asociality (items 5,6) and avolition (items 7,8) and BNSS Diminished expression was defined as the sum of the BNSS subscales blunted affect (items 9-11) and alogia (items 12,13); PANSS: Positive and Negative Syndrome Scale; HAMD: Hamilton Depression Rating Scale; CDSS: Calgary Depression Scale for Schizophrenia; GAF: Global Assessment of Functioning; PSP: Personal and Social Performance Scale. MWT IQ: Multiple Word Test Intelligence Quotient. ^1^Group differences were investigated using 2-sample t tests for continuous and *χ*^2^ tests for categorical data. For non-normally distributed data Mann-Whitney *U* tests were applied.

### Psychopathological data

To address our first aim, we compared the severity of NS in BD-I with SZ patients (Fig. 1 and Table 1 for statistics). Whereas the two groups did not differ in BNSS apathy (Fig. 1A), BD-I patients scored significantly lower on the BNSS expression subscale (Fig. 1B). Moreover, we observed a trend that BD-I patients have less total BNSS scores compared to SZ patients.

**Figure 1.**
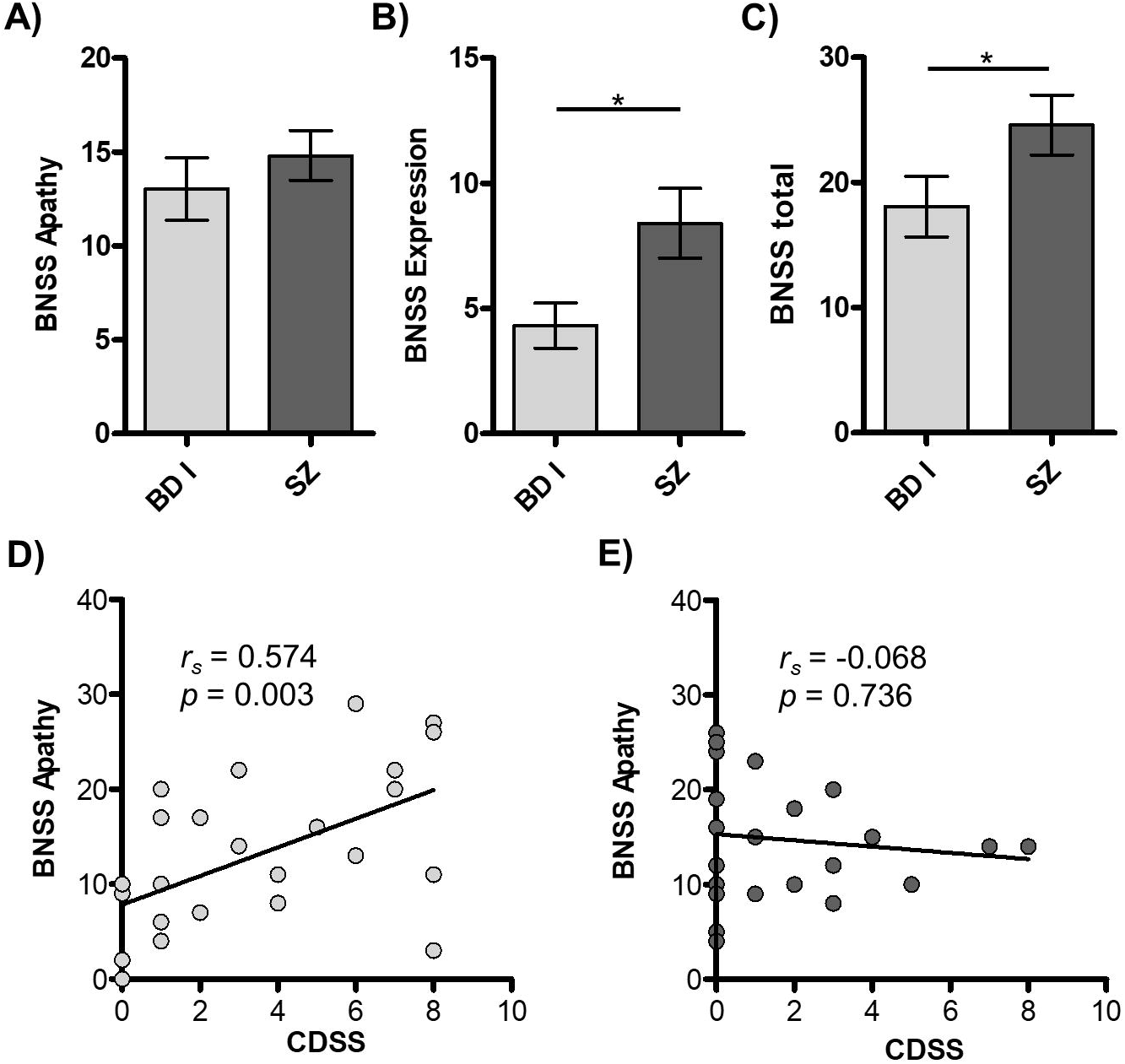
(A) Comparison of BNSS apathy between BD-I patients and patients with schizophrenia revealed no differences whereas (B) BD-I patients showed lower BNSS Expression and (C) higher CDSS total scores than patients with schizophrenia. There was a significant correlation between BNSS apathy and CDSS in (D) BD-I but not in (E) patients with schizophrenia. * = *p* < 0.05

BD-I patients showed higher CDSS scores than SZ patients (Fig. 1C). There was a significant correlation between BNSS apathy and CDSS in BD-I (Fig. 1D) but not in patients with schizophrenia (Fig. 1E). There was no significant correlation between BNSS diminished expression and CDSS in either group (BD-I: r_s_ = 0.268, *p* = 0.195; SZ: r_s_ = 0.174, *p* = 0.385). Given the significant correlation between apathy and subclinical depressive symptoms, we performed additional correlation analyses with the three subdomains anhedonia, avolition and asociality of the BNSS apathy factor. In BD, all three subdomains were correlated with depressive symptoms (anhedonia: rs = 0.599, p = 0.002; avolition: rs = 0.418, p =0.038; asociality: rs = 0.397 p = 0.049). Although these findings suggest a stronger association between anhedonia and depression, we did not observe a significant difference compared to the correlation between avolition and depression (Z = 1.60, p = 0.109) and the correlation between asociality and depression (Z = 1.464, p = 0.143). Additionally, the BNSS items comprising blunted affect did not correlate with the CDSS score in BD I (r_s_ = 0.294, p = 0.154) and SZ patients (r_s_ = 0.163, p = 0.418). In sum, these analyses suggest that in BD-I patients, subclinical depressive symptoms were associated with all three subdomains of the apathy factor.

Taken together, these findings revealed that BD-I patients have similar levels of apathy but lower levels of diminished expression compared to SZ patients. Interestingly, apathy was correlated with sub-syndromal depressive symptoms in BD-I patients but not in patients with SZ.

BD-I patients had lower scores on the PANSS positive symptom subscale than SZ patients. However, there was no correlation of positive symptoms with BNSS apathy in either of the groups (BD-I: r_s_ = – 0.019, *p* = 0.928; SZ: r_s_ = −0.136, *p* = 0.500). Regarding antipsychotic medication, chlorpromazine equivalents were higher in patients with SZ compared to BD-I patients (BD-I: 185.99 ± 259.8, SZ: 508.01 ± 369.2; U = 133, p < 0.001). However, there was no correlation with BNSS apathy in either of the groups (BD-I: r_s_ = 0.207, *p* = 0.320; SZ: r_s_ = −0.047, *p* = 0.816). Thus, no evidence for a contribution of positive symptoms or antipsychotic medication to the observed apathy symptoms was found.

## Behavioural data

### Comparison between BD-I patients and HC

The repeated-measures ANOVA with reward condition as within-subject factor and group affiliation as between-subject (BD-I patients and HC) factor revealed a significant main effect of reward (F(1.5,70) = 67.661, *p* < 0.001) but no significant effect of group (F(1, 47) = 2.196, p = 0.15) or group x reward interaction (F(1.5, 70) = 2.117, p = 0.14). Bonferroni *post-hoc* pairwise comparison of response times revealed significant differences between all reward conditions (all *p* < 0.001; no reward < low reward < high reward). These results indicate intact reward-related speeding in HC and BD-I. Furthermore, neither BNSS apathy nor BNSS expression correlated significantly with reward-related speeding (BNSS apathy: r = −0.062, p = 0.774; BNSS expression: r = 0.018, p = 0.933). Finally, we did not find group differences in total error rates (BD-I: 5.4 ± 3.6, HC: 5.7 ± 4.0; *U* = 293.5, *p =* 0.896) or total money gain (BD-I: 39.5 ± 4.5, HC: 38.9 ± 5.2; *U* = 274, *p* = 0.603) (Table S1).

### Comparison between BD-I patients and SZ patients

We repeated the same analysis to compare the BD-I patients with the previously published sample of chronic SZ patients (Kirschner *et al.*, 2016b). The repeated-measures ANOVA with reward condition as within-subject factor and group affiliation as between-subject factor (BD-I and SZ patients) revealed a significant main effect of reward (F(1.4, 67) = 39.376, *p* < 0.001) but no significant effect of group (F(1, 49) = 0.049, p = 0.83) or group x reward interaction effect was observed (F(1.4, 67) = 2.907, p = 0.08) (Table S1). None of the measures for task performance (response time, error rate, total gain) showed a significant correlation with IQ (estimated with the MWT) (Table S2).

## fMRI data

### Group comparison of VS and DS activation between BD-I patients and HC

A voxel-wise whole-brain analysis of the reward anticipation contrast in the combined sample of BD-I patients and HC revealed strong task-related activation in reward coding regions including the VS and DS (Table S6, Figure 2).

**Figure 2.**
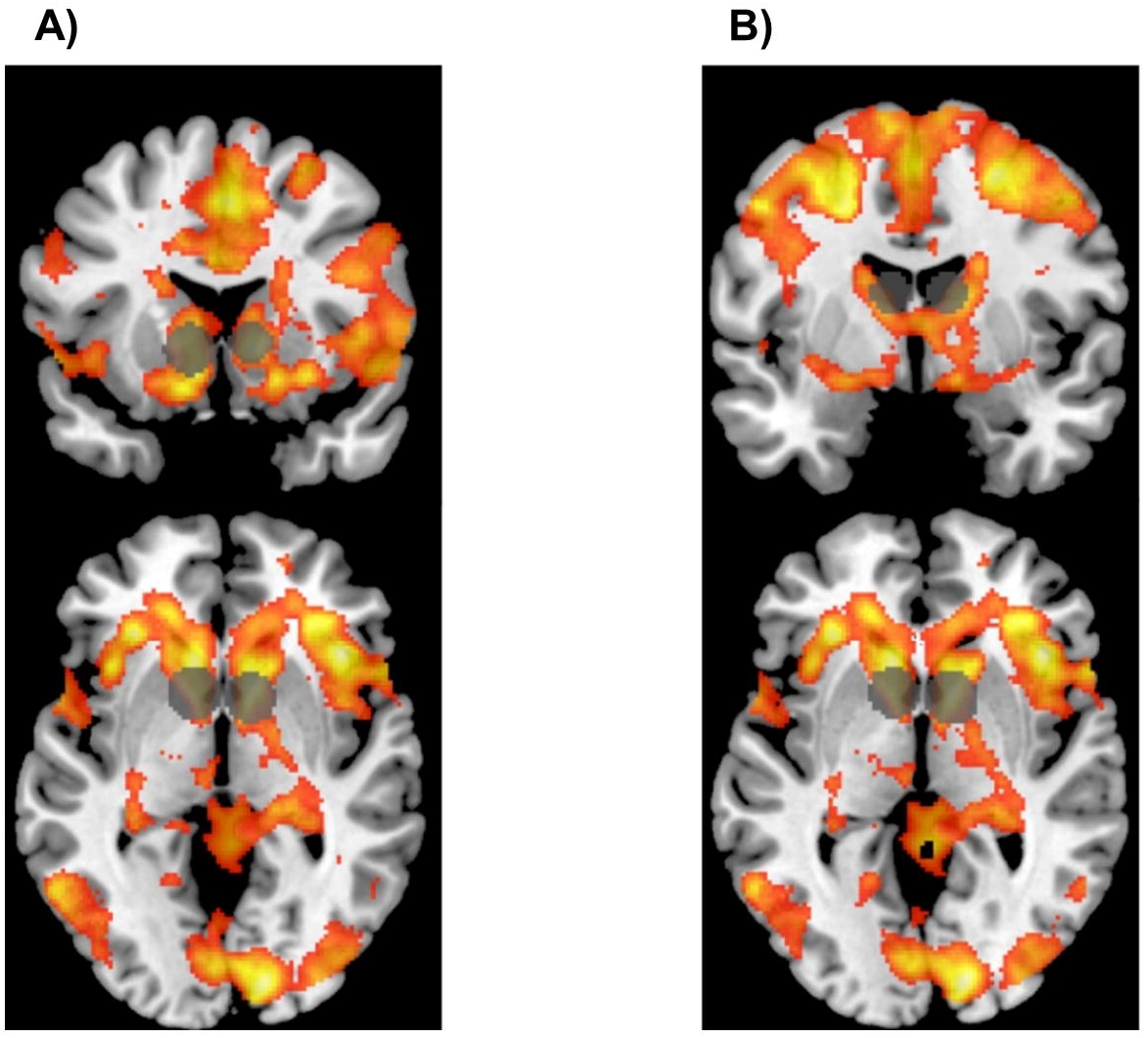
Activation map of the reward anticipation contrast (anticipation of high reward vs. anticipation of no reward) across the complete sample. Illustrated in coronal orientation (upper line) and axial (lower line) orientation. The VS ROI (A) and DS ROI (B) were overlaid on the activation map. The threshold was set at P<.05 FWE corrected.

To address the second aim of the study (relation of group to VS and DS activity), we compared the mean contrast signal from our a priori defined VS and DS ROI between HC and patients with BD-I (Figure 3A, C). There were no significant group differences in neural activation during reward anticipation in the VS (t = 0.412, p = 0.682) or DS (t = 1.09, p = 0.279). Thus, the VS and DS showed increased activity related to reward anticipation irrespective of group membership. Furthermore, in an explorative analysis we did not observe group differences in the low reward – no reward anticipation contrast and high reward – low reward anticipation contrast (Table S3).

**Figure 3.**
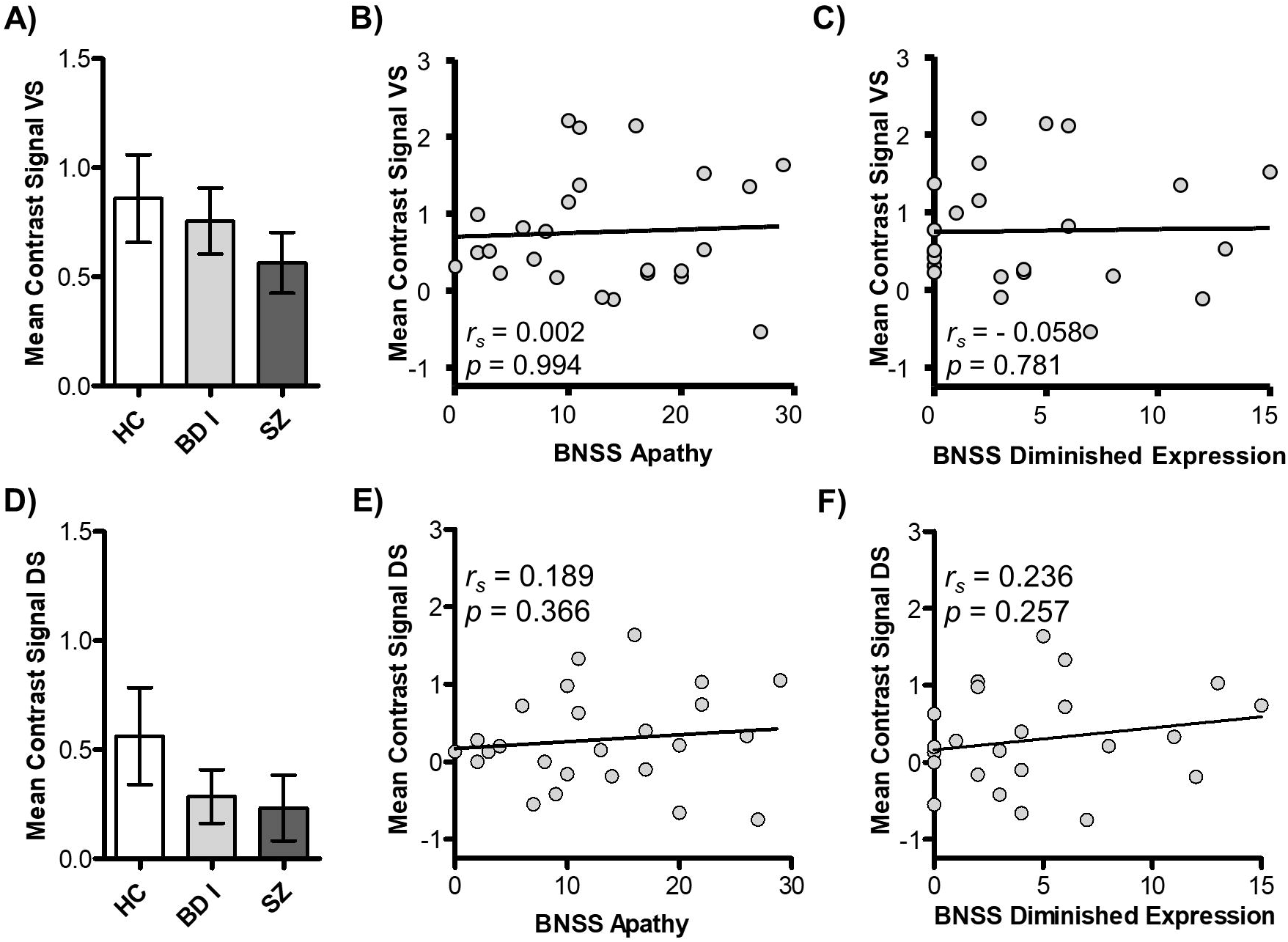
(A, D) Group comparison of activation during reward anticipation revealed no differences between HC and individuals with BD-I in VS (A) or DS (D). (B, C) VS activity during reward anticipation did not correlate with apathy or diminished expression. (E, F) DS activity during reward anticipation did not correlate with apathy or diminished expression.

### Group comparison of VS and DS activation between BD-I patients and SZ

In a next step we compared the mean contrast signal of the VS and DS ROI between BD-I patients and the previously published sample of chronic SZ patients (Kirschner *et al.*, 2016b). There were no significant group differences in neural activation during reward anticipation in the VS (t = 0.93, p = 0.357) or DS (t = 0.267, p = 0.79). Thus, VS and DS activations during reward anticipation did not differ significantly between BD-I patients and chronic SZ patients.

### Correlation analysis between reward anticipation and negative symptom domains

The third aim of the study was to investigate the association between the two negative symptom dimensions apathy and diminished expression and the mean contrast signal in our a priori ROIs of the VS and DS. Neither VS activity nor DS activity during reward anticipation were associated with one of the two negative symptom dimensions apathy or diminished expression (VS: apathy r_s_ = – 0.002, diminished expression r_s_ = – 0.058; DS: apathy r_s_ = 0.189, diminished expression r_s_ = 0.236, all p>.05) (Figure 3B, D). In sum, in patients with BD-I neither apathy nor diminished expression showed any association with reward anticipation in the striatum.

In a further analysis we investigated the potential correlation between depressive symptoms (CDSS Total score) and reward anticipation in the VS and DS. Again, we did not find any association between symptom expression and VS and DS activity (VS: r_s_ = 0.136, p = 0.515; DS: r_s_ = 0.145, p = 0.490). Furthermore, neither in BD-I patients nor in patients with SZ antipsychotic medication dose correlated with striatal activation during reward anticipation (SZ, VS: r_s_ = 0.128, p = 0.525, DS: r_s_ = 0.045, p = 0.824; BD-I, VS: r_s_ = 0.086, p =0.684, DS: r_s_ = −0.052, p = 0.804). Additionally, VS and DS activation during reward anticipation were not associated with functioning or education (Table S5). In the next step, we compared the correlation between apathy and VS activity in BD-I patients with the previous published negative correlation between apathy and VS activity in schizophrenia patients. Using Fisher r-to-z transformation we found a trend level significance (Z = 1.72, p = 0.085 two-tailed) for the differences between the two correlation coefficients suggesting that the association between apathy and ventral striatal activation was stronger in patients with schizophrenia than in patients with BD-I. Finally, we conducted explorative correlation analyses for the low reward – no reward anticipation contrast and high reward – low reward anticipation contrast, which revealed no significant correlation with apathy (Table S4).

### Extra-striatal dysfunction during reward anticipation

The fourth aim of the study was to identify potential extra-striatal prefrontal group differences in brain activity during reward anticipation. In our a priori IFG ROI, no group differences were observed comparing BD-I patients with HC and SZ patients. Furthermore, SZ patients did not differ from HC. Additional explorative whole brain analysis in BD-I patients revealed reduced activation during reward anticipation in the fusiform gyrus, lingual gyrus and precuneus compared to HC (Table S7).

In a next step, we aimed to identify potential associations between apathy and reduced activation in extra-striatal regions. Based on previous findings from our own group, we expected a negative association between apathy and IFG activation. Please, note that the main effect of high reward anticipation versus reward anticipation across the complete sample revealed activity in the right and left IFG supporting the relevance of this region in reward anticipation. In BD-I patients, we found a significant negative association between right IFG activation and apathy (56, 27, 9, cluster size = 41, t = 5.05, p=.034, peak-level FWE <0.05 corrected) (Fig 4). No association was observed in the left IFG. Please note that after Bonferroni correction for the number of ROIs this result was only at trend-level significance (p=.068). A secondary linear regression model with the log-transformed CDSS scores revealed no significant association between sub-syndromal depressive symptoms and reduced IFG activation. To test whether BD-I patients and SZ patients share the same apathy related IFG hypoactivation, we repeated the linear regression with apathy and IFG activation during reward anticipation in patients with SZ. In contrast to BD-I patients, we did not observe an association between apathy and reduced IFG activation in patients with SZ.

**Figure 4.**
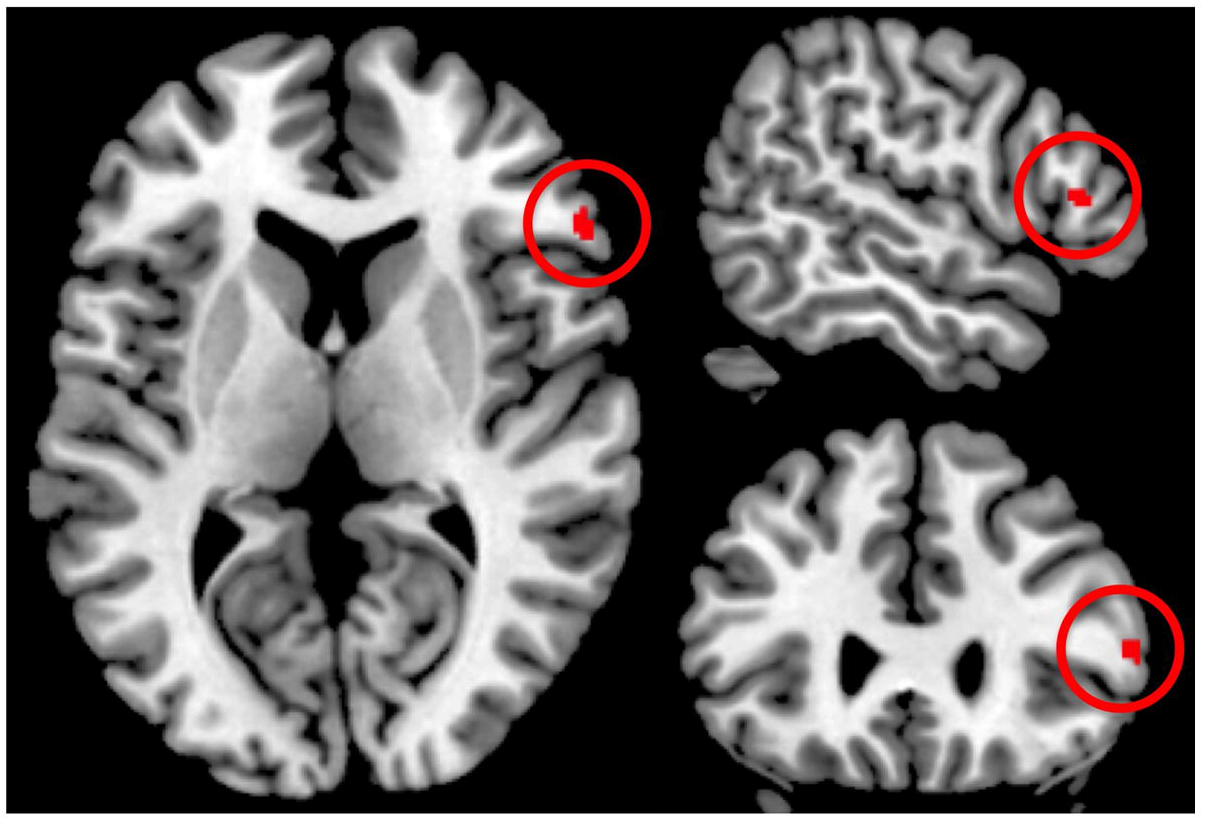
Associations between apathy and reduced activation in the right inferior frontal gyrus (56, 27, 9, cluster size = 41, t = 5.05, p=.034, peak-level FWE <0.05 corrected) in euthymic BD-I patients.

## Discussion

The present study investigated apathy and diminished expression in euthymic BD-I patients and its association with VS and DS activation during reward anticipation in a variant of the MID task. In euthymic BD-I patients, clinically measured severity of apathy was similar to patients with schizophrenia (Kirschner *et al.*, 2016b). However, BD-I patients showed less emotional expressivity deficits than patients with SZ. On the neural level, apathy was not associated with VS and DS activity during reward anticipation but showed an association with reduced prefrontal activation in the IFG.

It has recently been suggested that NS constitute a valid transdiagnostic psychopathological construct (Strauss *et al.*, 2016). In the present study, we confirm that NS can be assessed in euthymic BD-I patients and that the overall severity of both total BNSS and BNSS apathy is not different compared to a recently published study with schizophrenia patients (Kirschner *et al.*, 2016b). In contrast, BD-I patients scored less on the BNSS expression subscale than schizophrenia patients. Findings of similar scores on the BNSS apathy subscale but differences in BNSS expression are in line with a recent study by Strauss and colleagues (Strauss *et al.*, 2016). Although not fulfilling the criteria for a depressive episode, BD-I patients showed more depressive symptoms compared to SZ patients and these correlated significantly with BNSS apathy in the BD-I but not in the SZ group. Interestingly in euthymic BD-I patients, subclinical depressive symptoms were related to all three subdomains of the BNSS apathy factor: anhedonia, avolition and asociality. These findings suggest that when exploring NS in BD it might be difficult to differentiate symptoms such as anhedonia from other depressive symptoms (here assessed with the CDSS). This overlap and its potential impact on underlying neurobiological mechanism should be taken into account when investigating NS and depression in a transdiagnostic way. Furthermore, it is well known that BD-I patients in the euthymic state as determined by diagnostic criteria and structured interviews, display a broad range of neuropsychological and psychopathological impairments, including sub-syndromal depressive symptoms (Bourne *et al.*, 2013, Roux *et al.*, 2017). These residual depressive symptoms – in contrast to residual hypomanic symptoms – are an important predictor of functional outcome and quality of life (Bonnin *et al.*, 2010, Dias *et al.*, 2008).

When interpreting the results of the present study, it is therefore important to take into account the concept of secondary negative symptoms: Whereas primary negative symptoms are defined by being inherent to the disorder, secondary negative symptoms are thought to have different underlying sources such as depressive symptoms, productive psychotic features or medication (Kirkpatrick, 2014, Kirschner *et al.*, 2017). Our data clearly show that in euthymic patients with bipolar disorder motivational negative symptoms are at least in part secondary to sub-syndromal depressive symptoms. In contrast, we found no evidence for a contribution of other secondary sources of negative symptoms such as positive symptoms or medication side-effects. Taken together, the psychopathological findings of our study suggest that whereas NS can indeed be assessed as a transdiagnostic construct, the specific characteristics of the different subdomains are likely to differ between traditional categorical disease entities, emphasizing the importance of a detailed psychopathological assessment.

While the clinically measured severity of apathy in BD-1 patients was comparable to that of patients with schizophrenia, the specific association of apathy to sub-syndromal depressive symptoms in BD-I patients but not SZ patients, suggest different underlying neural mechanisms during reward anticipation: In contrast to the growing evidence that apathy is associated with reduced striatal activity in schizophrenia (Kirschner *et al.*, 2016b, Mucci *et al.*, 2015, Wolf *et al.*, 2014), this association was not observed in patients with euthymic BD-I. In addition, in line with previous studies in euthymic BD patients (Caseras *et al.*, 2013, Yip *et al.*, 2015) we did not observe an association between striatal activity and sub-syndromal depressive symptoms. Though speculative, one explanation for these divergent findings might be that primary and secondary negative symptoms have divergent neural correlates. While apathy, as a primary negative symptom, is associated with striatal dysfunction, apathy secondary to sub-syndromal depressive symptoms may be related to extra-striatal mechanisms. Our findings suggest that apathy in patients with BD-I might be more strongly related to cognitive processes relevant for action initiation and execution as reflected by prefrontal cortex activation (Barch and Dowd, 2010). Critically, the observed association between apathy and reduced IFG activation during reward anticipation is in line with previous results showing reduced IFG activation in bipolar depression (Redlich *et al.*, 2015). The relevance of prefrontal activity in euthymic and sub-syndromal depressive BD-I patients is underpinned by research reporting reduced top-down control of anteroventral prefrontal cortex on VS activity during reward processing (Trost *et al.*, 2014). Nevertheless, our findings have to be interpreted with caution given that only apathy but not depression was associated with reduced prefrontal cortex activation in the present study. In addition, some studies in bipolar depression, unipolar depression and schizophrenia have shown a correlation between depressive symptom scores and reduced striatal activity (Arrondo *et al.*, 2015, Hagele *et al.*, 2015, Satterthwaite *et al.*, 2015). Taken together, the potentially shared and distinct neural substrates of primary apathy and secondary apathy due to depression still have to be elucidated.

An additional explanation for these divergent findings might be the greater heterogeneity of neural activation during reward anticipation in different stages and subtypes of BD. While the meta-analysis of Radua and colleagues (Radua *et al.*, 2015) demonstrated that there is consistent evidence for reduced striatal activation during reward anticipation in SZ, the findings in BD are still inconclusive (Ashok *et al.,* 2017). Recently, Caseras and colleagues reported increased bilateral VS activity in BD-II patients, but no differences in BD-I patients compared to HC (Caseras *et al.*, 2013). This is in line with our observation of comparable activation during reward activation in euthymic BD-I patients and HC. However, a recent study by Yip et al. showed that during reward anticipation, unmedicated BD-II patients had decreased DS but not VS activity (Yip *et al.*, 2015). Here, similar to the literature on schizophrenia, which reported opposing results in unmedicated patients and patients treated with atypical antipsychotics (Nielsen *et al.*, 2012, Schlagenhauf *et al.*, 2008), the divergent findings might be due to differences in medication between the present study (medicated) and the previous report from Yip and colleagues (unmedicated) (Yip *et al.*, 2015). In sum, these findings point towards different mechanisms underlying motivational deficits between different stages and subtypes of BD. Of note, the explorative whole brain analyses of our study revealed reduced extra-striatal activation in fusiform gyrus, lingual gyrus and precuneus in BD-I compared to HC. These results replicated previous findings in BD-II patients (Yip *et al.*, 2015) and suggest that temporal, occipital and parietal regions involved in reward processing (Sescousse *et al.*, 2013) are impaired in euthymic BD-I patients.

### Limitations and future directions

There are several limitations to the present study: First, the sample size is only moderate and although we made an effort to recruit a homogeneous sample regarding the current disease stage, it is challenging to minimize variance in number of past depressive/manic episodes and duration of illness in chronic BD-I patients. These characteristics could mask specific mechanisms underlying various stages of the disorder. In particular, we did not use a conservative definition of euthymia (Samalin *et al.*, 2016), but allowed sub-syndromal depressive symptoms according to the definition of the ISBD Task Force (Tohen *et al.*, 2009). In order to elucidate specific neural correlates of different stages in BD, future neuroimaging studies should take advantage of precise recommendations to overcome the high variability in clinical criteria and duration for euthymia used in previous research (Robinson *et al.*, 2006, Samalin *et al.*, 2016, Tohen *et al.*, 2009). Furthermore, the present patient sample consists of medicated patients and does not take into account possible confounding effects of psychotropic drugs (Phillips *et al.*, 2008) since subgroups were too small to conduct *post hoc* comparisons between different drug classes. Third, the lack of specific measures for anhedonia other than the BNSS may have hampered the ability to identify distinct correlates for subdomains of apathy. Therefore, further studies should investigate detailed psychopathological characteristics including specific measures for anhedonia and motivational deficits during different episodes of the disorder. Especially, within-subject designs investigating different disease stages on an individual level could reveal important insights into the variability of the underlying neural mechanisms. Though speculative, understanding these processes could be an important step towards the development of targeted treatment interventions that address the oftentimes so burdensome switching between the different episodes of BD-I.

### Conclusion

The present study indicates that on a psychopathological level apathy is present in euthymic BD-I patients comparable to patients with schizophrenia but appear to be related to sub-syndromal depressive symptoms. This highlights the relevance of detailed clinical assessments of other symptom dimensions in transdiagnostic studies of apathy. The lack of convergent neural correlates of apathy in the striatum and the association of apathy with extra-striatal dysfunction in BD-I patients suggests divergent neurobiological mechanisms of apathy in BD-I patients compared to SZ patients. Thus, this transdiagnostic approach contributes to the empirical differentiation of shared and divergent pathophysiological mechanisms of apathy across psychiatric disorders. Taken together, these findings strengthen the notion to investigate motivational deficits in a dimensional approach as conzeptualized in the Research Domain Criteria (Cuthbert and Insel, 2010) in order to foster progress for new treatments (Strauss & Cohen 2017; Husain & Roiser 2018).

## Acknowledgments

This study was supported by the Swiss National Science Foundation (Grant No. 105314_140351 to Stefan Kaiser). Matthias Kirschner was supported by the National Bank Fellowship Award (Montreal Neurological Institute, McGill University). Philippe Tobler was supported by the Swiss National Science Foundation (PP00P1_150739, and PP00Pl_00014_l65884). We are grateful to Dr. Philipp Staempfli for his excellent technical support. Furthermore, we would like to thank all participants for their time and interest in our study.

## Ethical standards

The authors assert that all procedures contributing to this work comply with the ethical standards of the ethics committee of the canton of Zurich, Switzerland and with the Helsinki Declaration of 1975, as revised in 2008.

