## Supplement for "Shared and dissociable features of apathy and reward system dysfunction in bipolar I disorder and schizophrenia"

*Supplemental Information*

1. Supplementary Methods
   1.1. Neuropsychological Assessment
   1.2. Functional image acquisition
   1.3. Image pre-processing
   1.4. Fig. S1. Schematic illustration of the variant of Monetary Incentive Delay Task (MID)
   1.5. Fig. S2. Graphical illustration of pay-out structure
2. Supplementary Results
   2.1. Table S1. Behavioural results of the variant of the Monetary Incentive Delay Task
   2.2. Table S2. Correlation between MWT IQ and measures of task performance
   2.3. Table S3. VS and DS activation during low and high reward anticipation for HC and BD-1 patients
   2.4. Table S4. Spearman rank correlation between VS and DS activation and low reward anticipation and symptoms in BD-I patients
   2.5. Table S5. Correlation between VS and DS activation during reward anticipation and functioning and education
   2.6. Table S6. Whole brain analysis of reward anticipation across the complete sample
   2.7. Table S7. Whole brain group differences of reward anticipation
3. References
1. Supplementary Methods
   1.1. Neuropsychological Assessment

We assessed verbal learning (Auditory Verbal Learning Memory Test) (Helmstaedter C, Lendt M, Lux S. 2001), verbal and visual short-term working memory (Härting C, Markowitsch HJ, Neufeld H 2000), Corsi block-tapping (Kessels *et al.* 2000), processing speed (Digit-Symbol Coding) (Von Aster M, Neubauer A, Horn R. 2006), planning (Tower of London) (Shallice 1982) and semantic and phonetic fluency (animal naming, s-words) (Delis DC, Kaplan E, Kramer J. 2001). Results of all cognitive tests were summarized in a composite cognition score computed with the mean of z-transformed scores (based on HC group data). Additionally, we used the Multiple Word Test (Lehrl *et al.* 1995) to control for premorbid verbal intelligence.

1.2. Functional image acquisition

Imaging data were collected using a Philips Achieva 3.0 T magnetic resonance scanner with a 32-channel SENSE head coil at the MR Centre of the Psychiatric Hospital, University of Zurich. Functional MRI scans were acquired in 2 runs with 195 images in each run. A gradient-echo T2*weighted echo-planar image (EPI) sequence with 38 slices acquired in ascending order was used. Acquired in-plane resolution was 3 × 3 mm^2^, 3 mm slice thickness and 0.5 mm gap width over a field of view of 240 × 240 mm, repetition time 2000 ms, echo time 25 ms and flip angle 82°. The first 5 scans were discarded to eliminate the influence of T1 saturation effects. Slices were aligned with the anterior–posterior commissure. Anatomic data were acquired using an ultrafast gradient echo T1-weighted sequence in 160 sagittal plane slices of 240 × 240 mm resulting in 1 × 1 × 1 mm voxels.

1.3. Image pre-processing

Functional images were corrected for differences in the time of slice acquisition and motion using the realign and unwarp function of SPM8. We used a voxel displacement map, calculated from double phase and magnitude field map data, to correct for combined static and dynamic distortions. Furthermore, segmentation, bias correction and spatial normalization were performed. Finally, images were smoothed using a 6 mm full-width at half-maximum Gaussian kernel. To assure adequate quality of fMRI data, participants with translational head movement greater than 3 mm or extensive signal dropout in the EPI sequences were excluded (3 HC and 3 patients with SZ).

1.4. Figure S1. Schematic illustration of the variant of Monetary Incentive Delay Task (MID) adapted from Kirschner *et al.* 2015

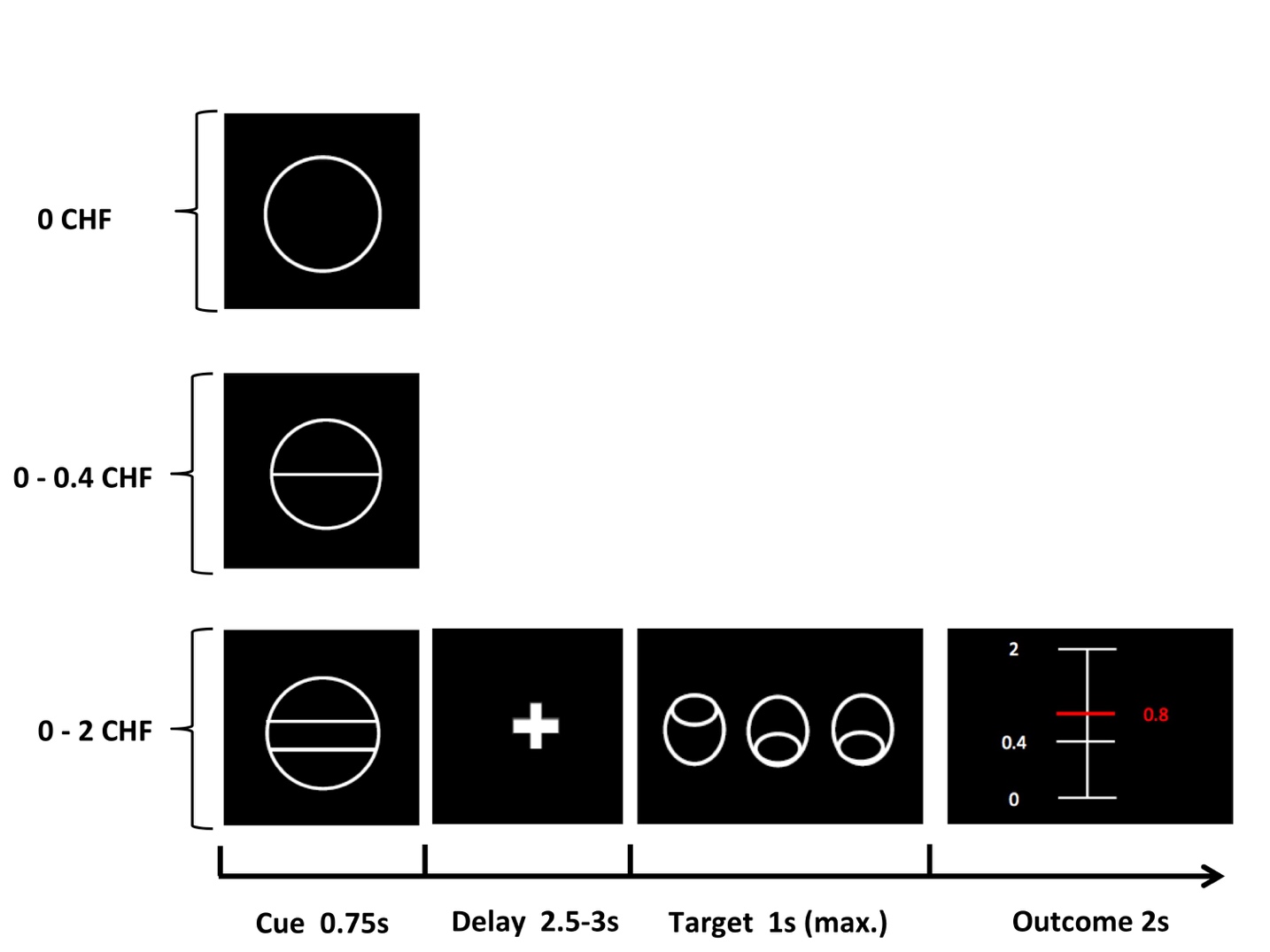

In each trial, participants saw 1 of 3 cues, indicating the amount of money to be won. After a delay period, participants had to identify an outlier from an array of 3 circles by pressing a correct button (either left or right). Immediately after a correct button press, participants were informed via visual feedback about the amount of money they had won during the current trial. A red horizontal line on the column ranging from the minimal amount (0 CHF) to the maximal win amount (2.0 CHF) indicated the precise amount of money won in each trial.

1.5. Figure S2. Graphical illustration of pay-out structure. Adapted from Simon *et al.* 2015

| payout % | time range |
| --- | --- |
| 100 | 2 |
| 99 | 1.95 |
| 98 | 1.9 |
| 97 | 1.85 |
| 96 | 1.8 |
| 95 | 1.75 |
| 94 | 1.7 |
| 93 | 1.65 |
| 92 | 1.6 |
| 91 | 1.55 |
| 90 | 1.5 |
| 89 | 1.45 |
| 88 | 1.4 |
| 87 | 1.35 |
| 86 | 1.3 |
| 85 | 1.25 |
| 84 | 1.2 |
| 83 | 1.15 |
| 82 | 1.1 |
| 81 | 1.05 |
| 80 | 1 |
| 79 | 0.95 |
| 78 | 0.9 |
| 77 | 0.85 |
| 76 | 0.8 |
| 75 | 0.75 |
| 74 | 0.7 |
| 73 | 0.65 |
| 72 | 0.6 |
| 71 | 0.55 |
| 70 | 0.5 |
| 69 | 0.45 |
| 68 | 0.4 |
| 67 | 0.35 |
| 66 | 0.3 |
| 65 | 0.25 |
| 64 | 0.2 |
| 63 | 0.15 |
| 62 | 0.1 |
| 61 | 0.05 |
| 60 | 0 |
| 59 | -0.0344 |
| 58 | -0.0688 |
| 57 | -0.1032 |
| 56 | -0.1376 |
| 55 | -0.172 |
| 54 | -0.2064 |
| 53 | -0.2408 |
| 52 | -0.2752 |
| 51 | -0.3096 |
| 50 | -0.344 |
| 49 | -0.3784 |
| 48 | -0.4128 |
| 47 | -0.4472 |
| 46 | -0.4816 |
| 45 | -0.516 |
| 44 | -0.5504 |
| 43 | -0.5848 |
| 42 | -0.6192 |
| 41 | -0.6536 |
| 40 | -0.688 |
| 39 | -0.7224 |
| 38 | -0.7568 |
| 37 | -0.7912 |
| 36 | -0.8256 |
| 35 | -0.86 |
| 34 | -0.8944 |
| 33 | -0.9288 |
| 32 | -0.9632 |
| 31 | -0.9976 |
| 30 | -1.02 |
| 29 | -1.05 |
| 28 | -1.1 |
| 27 | -1.15 |
| 26 | -1.2 |
| 25 | -1.25 |
| 24 | -1.3 |
| 23 | -1.35 |
| 22 | -1.4 |
| 21 | -1.45 |
| 20 | -1.5 |
| 19 | -1.55 |
| 18 | -1.6 |
| 17 | -1.65 |
| 16 | -1.7 |
| 15 | -1.75 |
| 14 | -1.8 |
| 13 | -1.85 |
| 12 | -1.9 |
| 11 | -1.95 |
| 10 | -2 |
| 9 | -2.05 |
| 8 | -2.1 |
| 7 | -2.15 |
| 6 | -2.2 |
| 5 | -2.25 |
| 4 | -2.3 |
| 3 | -2.35 |
| 2 | -2.4 |
| 1 | -2.45 |
| 0 | -2.5 |

Pay-out structure of the Monetary Incentive Delay task variant. For every individual, we calculated the 15 previous response times and sorted them from fast to slow using a simple bubble sorting procedure. We then selected the response times corresponding to the 60^th^ percentile and 80^th^ percentile, defining a minimum and maximum of the time range within each participant had to respond in order to win money (grey area = time range 0 to 1). Due to the fact that these ranges tended to be small (~5 to 10 milliseconds), we dispersed the time frame, ranging from -2.5 ranges below the original minimum and 1 range above the original maximum. Finally, the pay-out amount was determined by plotting the amount won in each trial with this modified dispersion of the original time range. With respect to the pay-out structure this approach gave the task a realistic feel by providing more dispersed outcome amounts. The X-axis represents the corresponding percentage of the maximal possible win during each trial. For example: in order to win 40% of the maximum amount during the CHF 2 condition (40%=CHF 0.8), participants had to respond at -1 range below the minimum. In order to win 80% (CHF 1.6), the response time had to be 1 range above the minimum.

2. Supplementary Results
   2.1. Table S1. Behavioural results of the variant of the Monetary Incentive Delay Task

|  | **BD-I**  **N=25** | **SZ**  **N=27** | **HC**  **N=25** | **BD-I vs. HC** | **BD-I vs. SZ** |
| --- | --- | --- | --- | --- | --- |
| Response time, ms | 37.3 ± 9.1 | 31.9 ± 7.4 | 33.1 ± 9.7 | reward: F(1.5,70) = 67.661, *p* < 0.001  group: F(1, 47) = 2.196, p = 0.15  group X reward: F(1.5, 70) = 2.117, p = 0.14 | reward: F(1.4, 67) = 39.376, *p* < 0.001  group: F(1, 49) = 0.049, p = 0.83  group X reward: F(1.4, 67) = 2.907, p = 0.08 |
| No reward | 566.8 ± 90.4 | 555.4± 111.0 | 519.2 ± 81.0 |  |  |
| Low reward | 518.4 ± 92.8 | 539.4 ± 106.1 | 490.3 ± 78.6 |  |  |
| High reward | 497.0 ± 89.3 | 505.3 ± 102.7 | 467.5 ± 80.6 |  |  |
| Error rate | 5.4 ± 3.6 | SZ: 6.3 ± 4.4 | 5.7 ± 4.0 | *U* = 293.5, *p* = 0.896 | *U* = 294.5, *p* = 0.576 |
| Total gain, CHF | 39.5 ± 4.5 | 36.1 ± 4.6 | 38.9 ± 5.2 | *U* = 274, *p* = 0.603 | *U* = 217, *p* = 0.043 |
| Notes: Data are presented as means and standard deviations. Potential group differences were investigated using 2-way repeated measures analysis of variance for response time and Mann-Whitney U tests for error rates and total gain. | | | | | |

2.2. Table S2. Correlation between MWT IQ and measures of task performance

|  | HC (n=25) | BD-I (n=23) | SZ (n=27) |
| --- | --- | --- | --- |
| No reward | rs = -0.215 | rs = 0.238 | rs = -0.062 |
| Low reward | rs = 0.11 | rs = 0.307 | rs = 0.269 |
| High reward | rs = -0.265 | rs = -0.347 | rs = 0.209 |
| Error rate | rs = -0.265 | rs = -0.347 | rs = 0.209 |
| Total gain, CHF | rs = 0.11 | rs = 0.307 | rs = 0.269 |
| Spearman rank correlation, all ps>0.1. | | | |

2.3. Table S3. VS and DS activation during low and high reward anticipation for HC and BD-1 patients

| (A) One sample t-test VS and DS activation during low and high reward anticipation in HC (n=25) | | | | |
| --- | --- | --- | --- | --- |
|  | Contrast Estimate | | Test statistic (T) | p-value |
| VS reward anticipation |  |  |  |  |
| LowRew - NoRew | 0.42 | (-0.64) | 3.30 | 0.003 |
| HighRew -LowRew | 0.43 | (-0.66) | 3.29 | 0.003 |
| HighRew - NoRew | 0.86 | (1.01) | 4.27 | <0.001 |
| DS reward anticipation |  |  |  |  |
| LowRew - NoRew | 0.10 | (0.89) | 0.55 | 0.589 |
| HighRew -LowRew | 0.46 | (0.71) | 3.27 | 0.003 |
| HighRew - NoRew | 0.56 | (1.11) | 2.54 | 0.018 |

| (B) One sample t-test VS and DS activation during low and high reward anticipation in BD-I patients (n=25) | | | | |
| --- | --- | --- | --- | --- |
|  | Contrast Estimate | | Test statistic (T) | p-value |
| VS reward anticipation |  |  |  |  |
| LowRew - NoRew | 0.36 | (0.52) | 0.36 | 0.002 |
| HighRew -LowRew | 0.39 | (0.48) | 0.39 | <0.001 |
| HighRew - NoRew | 0.76 | (0.75) | 0.76 | <0.001 |
| DS reward anticipation |  |  |  |  |
| LowRew - NoRew | -0.01 | (0.56) | -0.01 | 0.954 |
| HighRew -LowRew | 0.29 | (0.58) | 0.29 | 0.018 |
| HighRew - NoRew | 0.29 | (0.61) | 0.29 | 0.029 |

| (C) Group comparison VS and DS activation during low and high reward anticipation | | | | |
| --- | --- | --- | --- | --- |
|  | HC (n=25) | BD-I patient (n=25) | Test statistic (T) | p-value |
| VS reward anticipation |  |  |  |  |
| LowRew - NoRew | 0.42 (-0.64) | 0.36 (0.52) | 0.36 | 0.72 |
| HighRew -LowRew | 0.43 (-0.66) | 0.39 (0.48) | 0.25 | 0.80 |
| HighRew – NoRew | 0.86 (1.01) | 0.76 (0.75) | 0.41 | 0.68 |
| DS reward anticipation |  |  |  |  |
| LowRew - NoRew | 0.10 (0.89) | -0.01 (0.56) | 0.50 | 0.62 |
| HighRew -LowRew | 0.46 (0.71) | 0.29 (0.58) | 0.94 | 0.35 |
| HighRew - NoRew | 0.56 (1.11) | 0.29 (0.61) | 1.09 | 0.28 |

2.4. Table S4. Spearman rank correlation between VS and DS activation and low reward anticipation and symptoms in BD-I patients

|  | VS | | DS | |
| --- | --- | --- | --- | --- |
|  | LowRew - NoRew | HighRew -LowRew | LowRew - NoRew | HighRew -LowRew |
| Apathy (motivation and pleasure) | rs=-0.142 | rs=0.267 | rs=-0.209 | rs=0.332 |
| CDSS total score | rs=0.11 | rs=0.05 | rs=0.005 | rs=0.142 |
| all ps >0.1 * p=0.036 uncorrected. | | | | |

2.5. Table S5. Correlation between VS and DS activation during reward anticipation and functioning and education

|  | HC (n=25) | | BD-I (n=25) | | SZ (n=27) | |
| --- | --- | --- | --- | --- | --- | --- |
|  | VS | DS | VS | DS | VS | DS |
| *GAF score* | NA | NA | 0.008 | -0.203 | 0.204 | 0.18 |
| *PSP total score* | NA | NA | 0.084 | -0.109 | 0.296 | 0.169 |
| *Education (yr)* | 0.364 | 0.231 | 0.383 | 0.327 | -0.267 | -0.102 |
| Spearman rank correlation rs, all ps>0.05. | | | | | | |

To address, whether functioning (GAF, PSP) and education (number of education years) were associated with activity during reward anticipation we first run correlational analyses within each group separately. None of the three variables were significantly correlated with VS or DS activation during reward anticipation (see Table S5). Furthermore, we did not observed any group differences between BD-I patients and SZ patients when entering GAF score, PSP total score and education as covariate (VS (4,46) F = 0.396, p = 0.81; DS (4,46) F = 0.137, p = 0.968).

2.6. Table S6. Whole brain analysis of reward anticipation across the complete sample

|  | X | Y | Z (mm) | cluster size | T |
| --- | --- | --- | --- | --- | --- |
| Postcentral Gyrus | -40 | -19 | 51 | 2978 | 9.01 |
| Inferior Parietal Lobule | -45 | -30 | 46 |  | 8.63 |
| Precentral Gyrus | -39 | -22 | 61 |  | 7.85 |
| Medial Frontal Gyrus | -8 | 24 | -12 | 866 | 8.7 |
|  | -12 | 16 | -15 |  | 7.45 |
| Caudate | -10 | 21 | 0 |  | 7.41 |
| Fusiform Gyrus | -30 | -88 | -14 | 841 | 8.07 |
| Cerebellum | -18 | -81 | -18 |  | 7.6 |
| Lingual Gyrus | -12 | -88 | -12 |  | 7.1 |
| Cuneus | 14 | -93 | 4 | 829 | 7.65 |
| Inferior Occipital Gyrus | 15 | -90 | -9 |  | 7.11 |
| Middle Occipital Gyrus | 12 | -88 | 12 |  | 7.09 |
| Inferior Frontal Gyrus | 40 | 24 | -2 | 502 | 7.5 |
|  | 32 | 34 | 0 |  | 7.02 |
|  | 34 | 28 | -8 |  | 5.97 |
| Medial Frontal Gyrus | 8 | 27 | -9 | 706 | 7.47 |
| Caudate | 10 | 18 | -5 |  | 7.47 |
|  | 10 | 6 | 9 |  | 6.62 |
| Inferior Frontal Gyrus | -30 | 30 | -5 | 127 | 7.24 |
| Medial Frontal Gyrus | 2 | 17 | 43 | 893 | 7.14 |
|  | -2 | 3 | 57 |  | 7.03 |
|  | 3 | 11 | 52 |  | 6.52 |
| Middle Frontal Gyrus | 26 | -3 | 54 | 263 | 7.13 |
|  | 32 | -3 | 46 |  | 6.22 |
|  | 36 | 0 | 55 |  | 6.15 |
| Middle Frontal Gyrus | 33 | 33 | 28 | 95 | 7.07 |
| Precuneus | -12 | -67 | 46 | 367 | 6.97 |
|  | -4 | -72 | 49 |  | 6.66 |
|  | -6 | -66 | 57 |  | 6.3 |
| Cerebellum | 38 | -51 | -33 | 114 | 6.96 |
| Inferior Parietal Lobule | -34 | -51 | 54 | 166 | 6.8 |
|  | -27 | -49 | 51 |  | 6.47 |
| Superior Parietal Lobule | -21 | -55 | 57 |  | 5.69 |
|  | 12 | -67 | 48 | 129 | 6.71 |
|  | 15 | -60 | 48 |  | 6.07 |
| Middle Temporal Gyrus | -36 | -81 | 24 | 119 | 6.61 |
|  | -27 | -81 | 31 |  | 6.17 |
|  | -26 | -69 | 31 |  | 6.06 |
| Cerebellum | 8 | -75 | -23 | 103 | 6.48 |
|  | 15 | -82 | -21 |  | 6.16 |
| Caudate | 15 | -3 | 15 | 59 | 6.13 |
| Voxel-wise whole brain analysis of the contrast high reward anticipation > no reward anticipation, across all subjects, cluster-level FWE-corrected p<0.05, with a cluster-defining voxel-level threshold of *p*<0.001 uncorrected. | | | | | |

2.7. Table S7. Whole brain group differences of reward anticipation

| **a) HC > BD-I** | x | y | z | cluster size | T |
| --- | --- | --- | --- | --- | --- |
| Fusiform Gyrus^1^ | 46 | -46 | -11 | 183 | 5.01 |
| Fusiform Gyrus | -22 | -61 | -5 | 598 | 4.73 |
|  | -6 | -57 | -2 |  | 4.52 |
|  | -10 | -49 | -2 |  | 4.26 |
| Precentral Gyrus | 42 | -12 | 36 | 203 | 4.62 |
|  | 32 | -9 | 37 |  | 3.76 |
|  | 45 | -6 | 31 |  | 3.53 |
| Precuneus | -9 | -49 | 34 | 307 | 4.53 |
|  | -16 | -45 | 33 |  | 4.28 |
| Lingual Gyrus | -24 | -82 | 4 | 238 | 4.22 |
|  | -33 | -73 | 9 |  | 4.12 |
|  | -16 | -87 | 1 |  | 3.81 |
| Superior Parietal Lobule | -27 | -78 | 40 | 214 | 4.16 |
|  | -33 | -70 | 33 |  | 3.58 |
| Voxel-wise whole brain group comparison of the contrast high reward anticipation > no reward anticipation, cluster-level FWE-corrected p<0.05 with a cluster-defining voxel-level threshold of p<0.001 uncorrected.^1^cluster-level pFWE=0.09. | | | | | |
